# Targeting xCT-mediated glutamate release normalizes tumor angiogenesis in the brain

**DOI:** 10.1101/134924

**Authors:** Zheng Fan, Thomas Broggini, Eduard Yakubov, Tina Sehm, Sebastian Schürmann, Eric P. Meyer, Nevenka Dudvarski Stankovic, Mirko HH. Schmidt, Marco Stampanoni, Marcus A. Czabanka, Robert Nitsch, Michael Buchfelder, Oliver Friedrich, Ilker Y. Eyupoglu, Nicolai E. Savaskan

**Affiliations:** Department of Neurosurgery, Universitätsklinikum Erlangen, Friedrich-Alexander University of Erlangen – Nürnberg (FAU), D-91054 Erlangen, Germany; Department of Neurosurgery, Charité - Universitätsmedizin Berlin, D-10117 Berlin, Germany; Department of Neurosurgery, Paracelsus University Nürnberg, Germany; Institute of Medical Biotechnology, Department of Chemical and Biological Engineering, Friedrich-Alexander-University Erlangen-Nürnberg (FAU), Paul-Gordan-Str.3, 91052 Erlangen, Germany; Institute of Molecular Life Sciences, University of Zurich UZH, 8008 Zurich, Switzerland; Molecular SignalTransduction Laboratories, Institute for Microscopic Anatomy and Neurobiology, School of Medicine, Johannes Gutenberg University, D-55128 Mainz, Germany; Institute of Biomedical Engineering, ETH Zurich, 8092 Zurich, Switzerland & Swiss Light Source, Paul Scherrer Institut, Villigen, Switzerland; BiMECON Ent. Berlin, Germany

**Keywords:** malignant glioma, epilepsy, comorbidity, endothelial cell, glutamate receptor signaling, angiogenesis

## Abstract

Brain tumors are among the most malignant primary tumors, hallmarked by angiogenesis, neuronal destruction and brain swelling. Inhibition of the glutamate-cystein antiporter xCT (system x_c_^−^/SLC7A11) alleviates seizures, neuronal cell death and tumor-associated brain edema. Here we show enhanced tumor vessel growth and increased brain edema in xCT-expressing brain tumors. Furthermore, xCT-mediated glutamate impacts directly on endothelial cells in an N-methyl-D-aspartate receptor (NMDAR) dependent manner with intracellular Ca^2+^ release. Cerebral intravital microscopy revealed that xCT-driven tumor vessels are functional and display increased permeability. Endothelial-cell-specific NMDAR1 knockout mice (GRIN^iΔEC^) show suppressed endothelial sprouting and vascular density compared to control littermates. In addition, implanted gliomas in GRIN^iΔEC^ mice display reduced tumor vessels in contrast to gliomas in wildtype animals. Moreover, therapeutic targeting of xCT in gliomas alleviates tumor angiogenesis to normalized levels comparable to controls. Our data reveal that xCT and its substrate glutamate specifically operate on endothelial cells and promote neoangiogenesis. Thus, targeting xCT expression and glutamate secretion in gliomas provides a novel therapeutic roadmap for normalizing tumor angiogenesis.

## INTRODUCTION

Primary malignant brain tumors (gliomas) are one of the deadliest neoplasia, which carry poor prognosis for patients despite current aggressive multimodal therapies (1); (2). Human gliomas are highly vascularized (3) and grow in a space-occupying manner and thereby impinge on eloquent (functional) brain structures causing profound neurological impairments. This process is often exacerbated by the mass effects of tumor-associated brain swelling leading to pressure-induced brain damage due to the limited space within the rigid skull. Brain edema crucially contributes to the clinical course and outcome of patients with gliomas (4) and has been confirmed in experimental settings with mice (5). One critical cytotoxic mechanism for edema formation has recently been identified with the glutamate transporter xCT at center stage (6). xCT (also known as SLC7A11 as the regulatory part of system x_c_^−^) is the catalytic domain of a heteromeric transporter including a heavy chain subunit (CD98/SLC3A2/4F2hc). Expression of both subunits is increased in patients suffering from malignant gliomas (7) (8). xCT imports cystine in exchange of the amino acid and neurotransmitter glutamate, which released into the extracellular milieu at neurotoxic concentrations (9) (10). Preclinical studies demonstrated that xCT inhibition in brain tumors reduces neuronal degeneration, glutamate-induced epilepsy and alleviates perifocal edema formation (6) (11). In addition, xCT expression correlates with malignancy in humans (7, 12). However, whether the glutamate released by the antiporter xCT additionally executes vasogenic functions which lead to brain edema is currently not defined (13). Also, whether glutamate receptor activation occurs in endothelial cells in relation to angiogenesis has not been investigated.

The glutamate specific N-methyl-D-aspartate receptors (NMDARs) are expressed in neurons governing neuronal plasticity and antioxidative defense in the CNS (Papadia et al., 2008; Baxter et al., 2015). NMDARs have been implicated in various diseases and neoplasia (Hardingham & Bading, 2010; Li & Hanahan, 2013). The NMDAR subunits NR1 NR2b in particular are associated with tumor promoting activity exhibiting receptor functionality in tumor cells (14).

To evaluate the influence of xCT-induced and glutamate on brain tumor growth apart from edema, we investigated the vasculature found in gliomas and assessed xCT-dependent tumor angiogenesis. These analyses allowed us to unravel the role of excitatory amino acid glutamate in tumor vessels and angiogenesis. We found that endothelial cells utilize NMDAR for vasculature formation, indicating a novel biological aspect of glutamate signaling.

## RESULTS

### Perturbed xCT expression impacts tumor angiogenesis

We first investigated whether alteration of the glutamate antiporter xCT would affect glioma-induced cell death in the tumor microenvironment. To gain detailed insights into this, we implanted GFP-expressing glioma cells (WT) into brain tissue harboring the complex microenvironment including all cellular constituents of the brain. Wild-type gliomas strongly proliferated in brain parenchyma, and this massive growth was accompanied by increased neuronal cell death (Suppl. Fig. 1 a, b). Interestingly, tumor-induced cell death was alleviated in xCT knock down (xCT^KD^) gliomas in comparison to wild-type tumors (81 ± 4% vs. 100 ± 5%) (Suppl. Fig. 1 a, b). To validate this finding, we analyzed gliomas overexpressing the glutamate transporter xCT (xCT^OE^ gliomas). These tumors showed increased levels of glutamate secretion into the extracellular environment compared to wild-type gliomas (69 ± 4 µM vs. 51 ± 3 µM Glu_ex_ in conditioned media), whereas xCT silenced gliomas showed diminished glutamate secretion (30 ± 5 µM Glu_ex_) (Suppl. Fig. 1 c). In addition, xCT overexpressing gliomas induced elevated levels of neuronal cell death exceeding the impact of wild-type glioma levels (135 ± 8% vs. 100 ± 5%) (Suppl. Fig. 1). Besides tumor-associated neurodegeneration, another relevant malignancy criterion in human gliomas is the extent of newly formed vessels, or tumor angiogenesis. Thus, we further analyzed the potency to induce angiogenesis in these tumors *ex vivo*. Wild-type rodent and human gliomas revealed augmented peritumoral vessel growth whereas xCT^KD^ gliomas showed normalized vessels with decreased numbers (Fig. 1 a, b; Suppl. Fig. 2). Interestingly, xCT^OE^ gliomas displayed increased radial vessels (Fig. 1 a, b; Suppl. Fig. 2). These data indicate that the glutamate transporter xCT in tumor cells is able to trigger neoangiogenesis and shapes tumor vessels.

**FIGURE 1:**
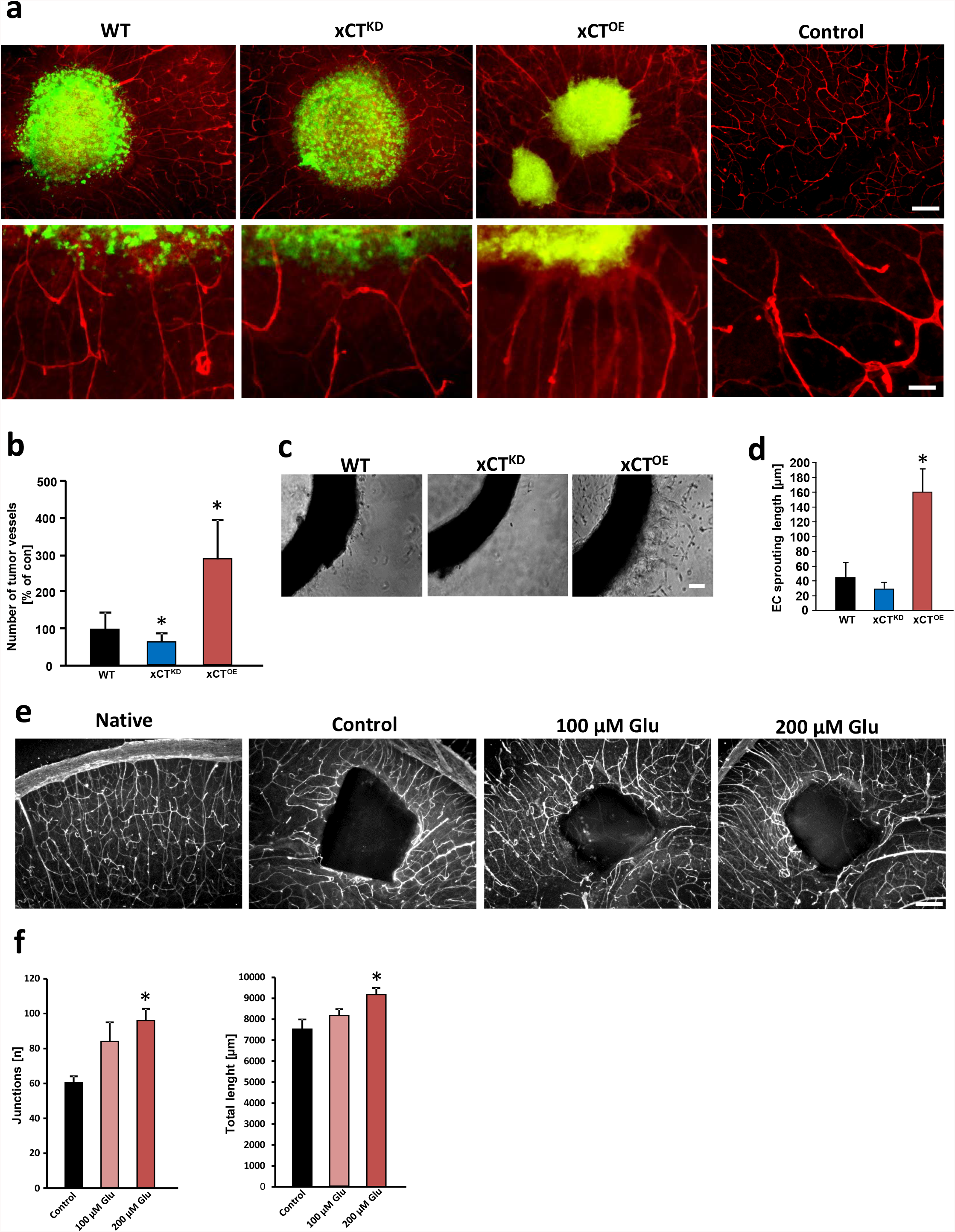
RNA-mediated xCT knockdown normalizes tumor angiogenesis ex vivo. **a**, RNAi-mediated xCT silencing (xCT^KD^) alleviates tumor-induced angiogenesis and xCT overexpressing tumors (xCT^OE^) show increased number of vessels. Upper, representative images from scrambled siRNA expressing GFP^+^ gliomas (WT), xCT siRNA knock down (xCT^KD^) and xCT overexpressing (xCT^OE^) gliomas 7 days after tumor implantation. The tumor is revealed by GFP expression (green) and vessels are immuno-stained for laminin (red). Scale bar represents 200 µm. Bottom, higher magnification of vessels within the peritumor area. Radial tumor vessel number in peritumor area is significantly lower in xCT^KD^ gliomas compared to WT-gliomas. xCT^OE^ gliomas show increased tumor vessel densities with abnormal inward directed and radial vessel structures. Scale bar at bottom represents 50 µm. **b**, Quantification of radial tumor vessels in peritumor areas of WT, xCT^KD^ and xCT^OE^ gliomas. Analysis was performed by manually counting the number of vessels which grew into the tumor divided by the tumor border length. Control group was set as 100%. Statistical significance was calculated (means are given as percentage of controls ± s.d., *P < 0.05, from n ≥ 6). **c**, Aortic explant cultures treated with conditioned media from wild-type glioma cells expressing scrambled siRNA (WT), xCT siRNA (xCT^KD^) and xCT overexpressing (xCT^OE^) gliomas. Representative images of aortic rings show endothelial sprouts which are denser and longer in explants treated with conditioned media from xCT^OE^ gliomas compared to WT gliomas, while in the conditioned medium of xCT^KD^ glioma cells, aortic rings show reduced outgrowth. Scale bar represents 100 µm. **d**, Quantification of endothelial outgrowth length. Statistical significance was calculated (means ± s.d., *P < 0.05, from n ≥ 4). **e**, Glutamate promotes vessel growth in native brain tissue. Representative images from native brain slices cultured under control conditions alone (native), in the presence of a PBS loaded sponge without glutamate (control), with low glutamate loaded sponge (100 µM Glu) and high glutamate loaded sponge (200µM Glu). Vessels are visualized with anti-laminin immunostaining (white signal). Scale bar represents 200 µm. **f**, Quantification of sponge assay with vessel junctions (left) and total vessel length (right). Statistical significance was calculated (means ± s.d., *P < 0.05, from n ≥ 4). All significance analyses were conducted using student’s t. test.

To gain additional insights into the mechanisms of xCT-induced angiogenesis, we investigated the effects of secreted factors derived from gliomas in established angiogenesis assays. For this, we applied conditioned media from wild-type, xCT^KD^ and xCT^OE^ gliomas to cultured aortic rings. Conditioned media derived from wild-type gliomas induced endothelial sprouting in cultured aortic explants whereas conditioned media from xCT^KD^ provoked solely minor effects (Fig. 1 c, d). In contrast, xCT^OE^ conditioned media promoted endothelial sprouting (Fig. 1 c, d). Since xCT^OE^ gliomas release high amounts of glutamate into the extracellular microenvironment, we speculated that this amino acid mediates the observed pro-angiogenic response. Therefore, we performed an organotypic brain vascular sponge assay and implanted glutamate loaded cubes onto brain sections and monitored the vascular response (Fig. 1 e). Glutamate-loaded cubes altered the vasculature dramatically and attracted many vessels in comparison to control cubes loaded solely with the solvent (Fig. 1 e, f).

We next cultured matrigel-embedded aortic explants in DMEM medium without supplements. After initial endothelial cell sprouting, we started to treat the tissue samples with glutamate and quantified the angiogenic response (Fig. 2 a, b). Treatment of aortic explant cultures with 100 µM glutamate increased endothelial sprouting significantly compared to controls (Fig. 2 a, b). To test whether this effect was caused by glutamate receptor-mediated signaling, we utilized various ionotropic glutamate receptor antagonists specifically directed against NMDA and AMPA subtypes. NMDARs were blocked with the non-competitive antagonist MK801 and we monitored the vascular response. The glutamate-induced endothelial sprouting effect was abolished when aortic rings were treated with the specific NMDAR antagonist MK801 (MK, 100 µM), whereas MK801 alone did not affect endothelial cell viability nor endothelial sprouting (Fig. 2 a, b). Next, we used the non-competitive AMPA-specific antagonist GYKI52466 (GYKI) applied at 50 µM to aortic explant cultures. In contrast, blocking glutamatergic AMPA receptors with GYKI did not abolish the outgrowth-promoting effects of glutamate. Endothelial sprouting was still significantly increased compared to controls (Fig. 2 b). These experiments indicate that glutamate induces endothelial sprouts probably specifically via NMDAR subtype signaling.

**FIGURE 2:**
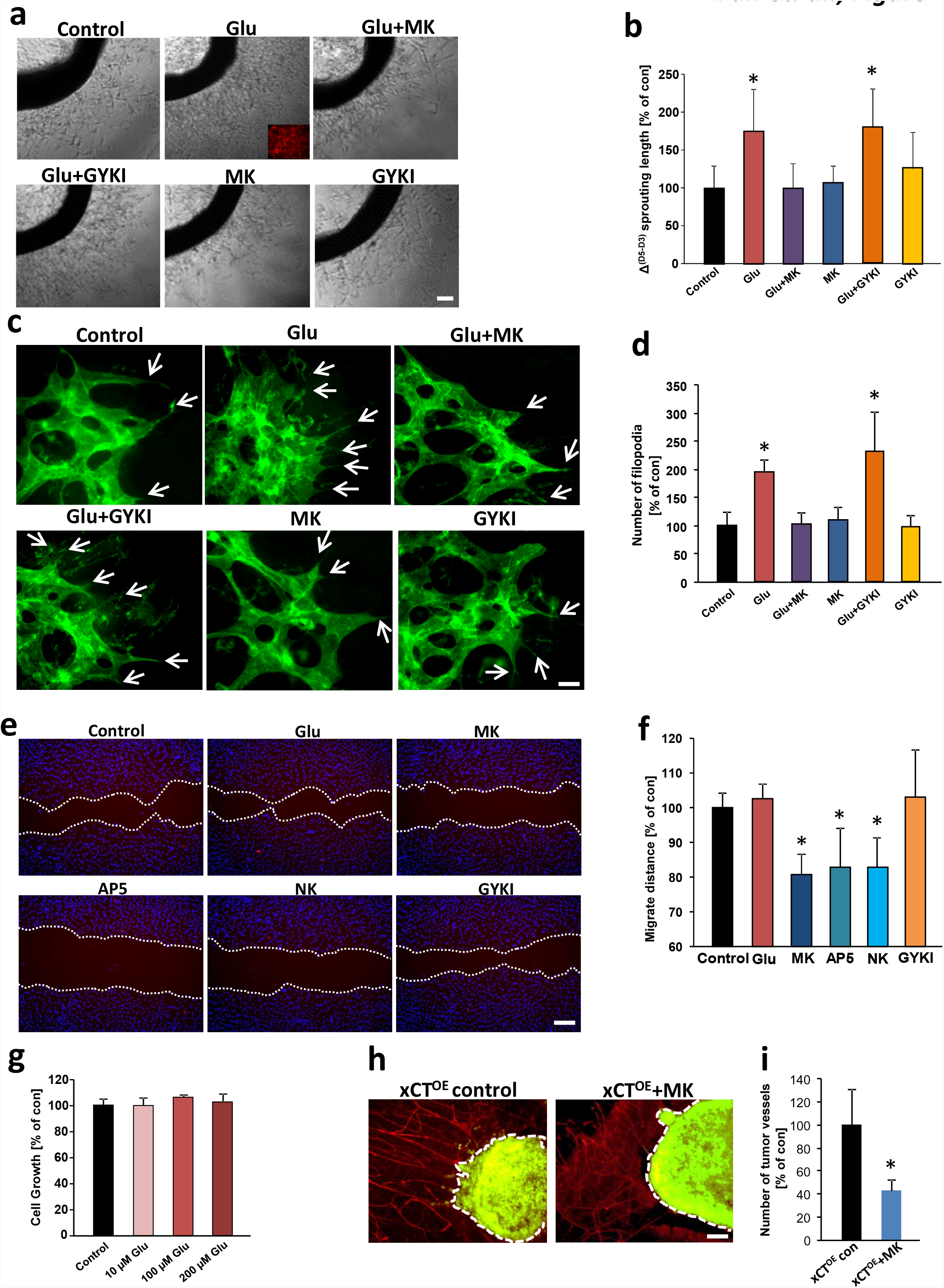
Glioma-derived glutamate promotes endothelial sprouting and angiogenesis. **a**, Glutamate promotes endothelial sprouting in aortic explant cultures. Aortic explants were treated with 100 µM glutamate alone, with 100 µM glutamate plus glutamate receptor antagonists MK801 (MK, 100 µM) or GYKI (50 µM). Representative images of aortic rings display that glutamate-treated aortic explants show stronger endothelial sprouting and vessel density compared to the control group. The glutamate-induced sprouting is inhibited in the presence of NMDA receptor antagonist MK801, while in the presence of the AMPA receptor antagonist GYKI, the glutamate-induced sprouting is not blocked. MK801 or GYKI alone did not show any toxic effects on endothelial sprouting compared to controls. Scale bar, 100 µm. **b**, Quantification of endothelial cell sprouting. Differences were shown in sprouting length (Δ of sprouting length) between day 5 and day 3 (start treatment). Δ sprouting length in control group was set as 100%. Statistical significance was calculated with one way ANOVA (means ± s.d., *P < 0.05, from n = 8). **c**, Representative images of isolectin-B4-stained angiogenic front of retinas treated with 50 µM glutamate (Glu), 50 µM glutamate plus 100 µM MK801 (Glu + MK), 50 µM glutamate plus 50 µM GYKI (Glu + GYKI), 50 µM GYKI alone (GYKI) and 100 µM MK801 alone (MK). Note the strongly enhanced angiogenesis in glutamate-treated retinas, white arrows indicate the filopodia. Scale bar represents 50 µm. **d**, Quantitative analysis of filopodia number per retina front length, control group was set to 100%. Data are given as mean ± s.d. (n = 4 per group) and statistically analyzed with one way ANOVA, *P < 0.05. **e**, NMDA receptor antagonists MK, AP5, and NK inhibit endothelial cell migration. Endothelial cell migration was monitored and endothelial cells were treated with 50 µM glutamate (Glu), NMDA receptor antagonist MK (100 µM MK801), 100 µM AP5 (AP5), 20 µM Norketamine (NK) and AMPA receptor antagonist (50 µM GYKI). Migration distance was measured following 12 hours of incubation. **f**, Quantification of endothelial cell migration after treatment for 12 hours. Values of control group were set to 100%. Statistical significance was calculated with one way ANOVA (means ± s.d., *P < 0.05, from n = 4). **g**, Monitoring human umbilical vein endothelial cell (HUVEC) proliferation following glutamate stimulation. HUVECs were treated with various concentrations of glutamate (10, 100 and 200 µM) for 48 hours and cell growth was assessed by MTT assay. Control group was set to 100%. n = 12 per group). **h**, Pharmacological inhibition of NMDA receptors antagonize xCT^OE^-induced angiogenesis. Representative images of glioma-implanted (green) brain slice cultures are given from untreated control group (xCT^OE^ control) and MK801 treated slices (xCT^OE^ + MK). Seven days after tumor implantation, tumor vessels were evaluated by laminin immunostaining (red). Note that pathological vessels’ radial pattern were diminished after MK801 treatment compared to control brain slices. **i**, Quantification of tumor vessel numbers/ tumor border length in xCT^OE^ gliomas under control condition and after MK801 treatment. Control group was set as 100%. Statistical significance was calculated with student’s t. test, n = 10 per group.

To redefine our observation for gliomas, rat brain endothelial cells were tested for their capacity to form vascular tubes under glutamate application. These experiments revealed that brain endothelial cells display increased total length of vascular tubes when treated with 100 µM glutamate (Suppl. Fig. 3). This glutamate-induced tube formation activity could be blocked with the NMDAR antagonist MK801 but remained intact with the AMPAR antagonists GYKI (Suppl. Fig. 3 a, b).

Next, we studied the role of glutamate *ex vivo* and monitored the retinal vasculature. Glutamate application significantly increased endothelial sprouts. This effect was reversed by simultaneous application of the NMDAR antagonist MK801, whereas inhibition of AMPA receptor signaling did not alter tube formation (Fig. 2 c, d). Next, we investigated the impact of glutamate signaling on endothelial cell migration. For this, we measured the migration distance after glutamate treatment and NMDA and AMPA receptor inhibition (Fig. 2 e). Additional glutamate application increased endothelial cell migration slightly in the endothelial growth medium with a basal glutamate level of 100 µM (Fig. 2 e, f). Moreover, NMDAR specific antagonists MK801, AP5, and NK reduced endothelial migration significantly, whereas AMPAR specific antagonists such as GYKI did not affect endothelial migration (Fig. 2 e, f).

To further analyze whether glutamate acts as an endothelial growth factor, we determined endothelial proliferation. Dose-response analysis revealed that glutamate at various concentrations ranging from 10 µM up to 200 µM does not significantly foster endothelial cell proliferation (Fig. 2 g). We corroborated these findings by testing whether interference with the NMDAR signaling can normalize xCT-induced tumor angiogenesis, i.e. shifting vascular parameters to control levels. For this, we compared brain tumor vessels under control conditions and MK801 application in xCT overexpressing gliomas. Noteworthy, blocking NMDAR in the organotypic tumor microenvironment (VOGiM) led to reduced numbers of tumor vessels compared to untreated controls (Fig. 2 h, i). Thus, these data indicate that glutamate acts directly on endothelial cells in an NMDAR-dependent manner.

### xCT-induced vessels are functionally perfused *in vivo*

We continued analyzing the vascular architecture in WT, xCT^KD^ and xCT^OE^ glioma implanted animals *in vivo* (Fig. 3). In contrast to WT gliomas, xCT^KD^ gliomas showed reduced vessel density (Fig. 3 a). Conversely, xCT^OE^ gliomas displayed an increased number of vessels (Fig. 3 a). Moreover, xCT^OE^ gliomas showed reduced neuronal cell survival (Fig. 3 b). Thus, these data corroborate the findings from the *ex vivo* VOGiM experiments indicating that xCT is an effective target for inhibiting tumor angiogenesis.

**FIGURE 3:**
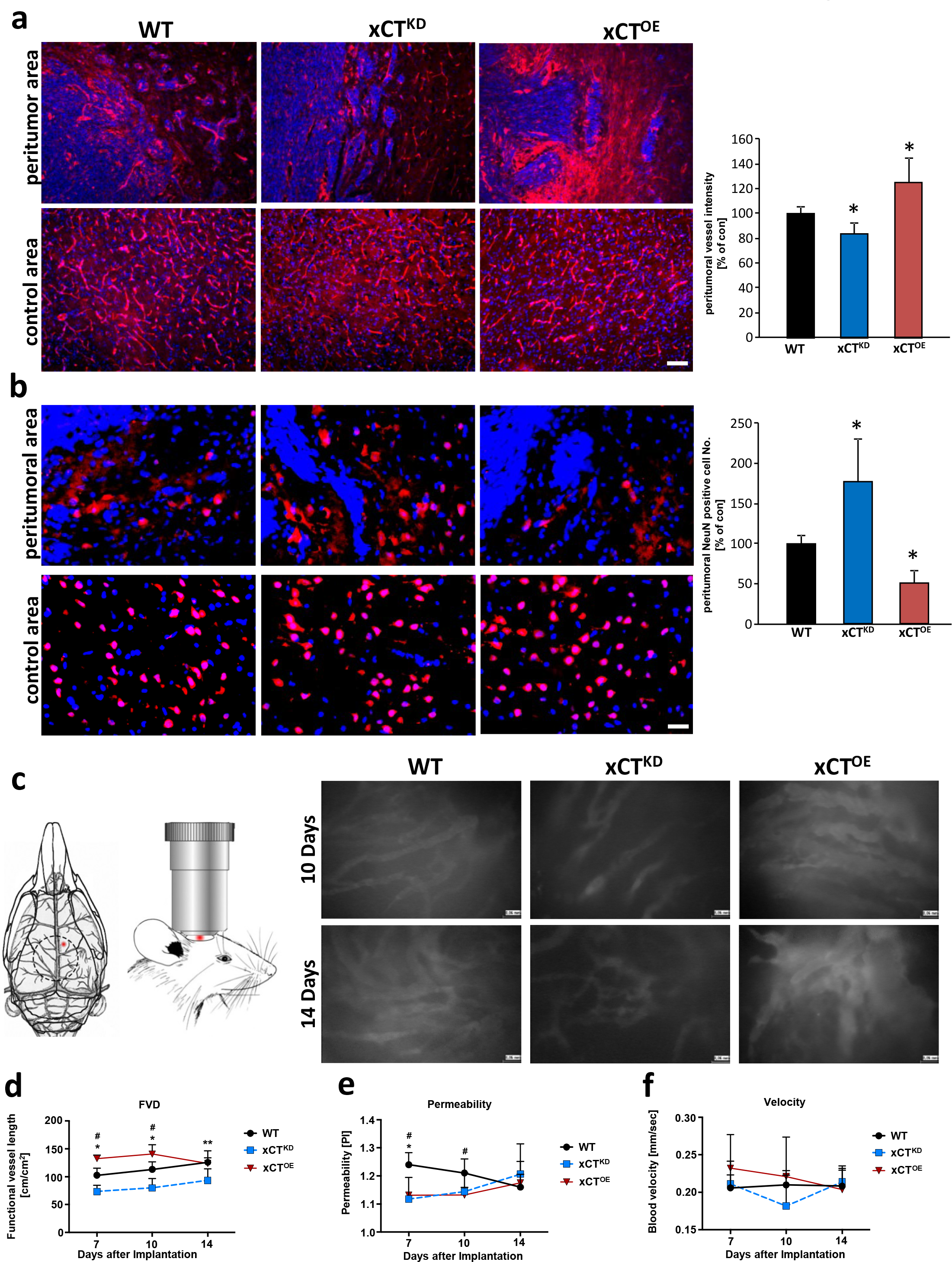
xCT-expressing tumors form functionally perfused vessels *in vivo*. **a**, *Left*, Representative immunofluorescence images for laminin (red) in the peritumor area (top row) and control area (bottom row) from wild-type gliomas (WT), xCT^KD^ gliomas (xCT^KD^), and xCT^OE^ gliomas (xCT^OE^). Wild-type tumor group is given as 100%. Nuclei are shown in blue. Scale bar represents 100 µm. *Right*, Quantification of peritumor vessel density in wild-type gliomas (WT), xCT^KD^ gliomas (xCT^KD^), and xCT^OE^ gliomas. Statistical significance was calculated with Students t-test, (means ± s.d., *P < 0.05, from n ≥ 4). **b**, *Left*, Neurodegeneration monitored in brains of wild-type gliomas (WT), xCT^KD^ gliomas (xCT^KD^), and xCT^OE^ gliomas (xCT^OE^). Representative images from NeuN immunostained (red) sections in peritumor areas (top) and control areas. Scale bar represents 50 µm. *Right*, Quantification of neurodegeneration in wild-type gliomas (WT), xCT^KD^ gliomas (xCT^KD^), and xCT^OE^ gliomas. Values of the WT glioma group is set as 100%. Statistical significance was calculated with student’s t. test, (means ± s.d., *P < 0.05, from n ≥ 4). **c**, Experimental set-up for monitoring vessel functionality. *Left*, Overview of murine skull and placement of cranial window. Tumor location is displayed by red dot. *Right*, Intravital microscopic images show typical tumor vasculature in control tumors (WT), low vascular density in xCT^KD^ gliomas (xCT^KD^), and dense and heterogeneous vascular morphology in xCT^OE^ gliomas (xCT^OE^). Scale bar represents 60 µm. **d-f**, Quantitative analysis of blood-brain barrier function of tumor vessels at day 7, 10 and 14 after spheroid tumor implantation. #: P < 0.05 xCT^OE^ vs. WT gliomas. *: P < 0.05 xCT^KD^ vs. WT tumors. Functional vessel density (FVD), permeability index (PI, intravascular to extravascular fluorescence intensity ratio) and velocity are given in wild-type gliomas (WT), xCT^KD^ gliomas (xCT^KD^), and xCT^OE^ gliomas. (n ≥ 5 per group).

Next, we investigated the functionality of newly formed vessels to estimate the involvement of xCT in vasogenic edema. For this, we performed cerebral intravital microscopy on mice bearing orthotopically implanted brain tumor spheroids. Interestingly, brain tumors from xCT^OE^ glioma cells showed increased values of functional vessel length compared to wild type gliomas (Fig. 3 c-f, significant values marked with hash). Hence, xCT^OE^ gliomas displayed high microvascular permeability at day 7 and 10 compared to wild-type gliomas and approached similar permeability values as wild-type gliomas at later time points (Fig. 3 c-f). Conversely, xCT^KD^ gliomas showed reduced functional vessel density whereas the microvascular permeability and velocity were comparable to values of wild-type gliomas (Fig. 3 d-f, significant values marked with star). Thus, xCT^OE^ gliomas promote tumor vessel formations which are perfused and functional for nurturing tumor growth.

### NMDAR mediated glutamate signaling in endothelial cells

To unveil the underlying mechanisms of glutamate mediated-effects on the tumor vasculature, we tested the endothelium for NMDAR expression. First, we analyzed the expression of the NMDAR subtypes in whole aortic tissue to confirm the responsiveness to glutamate. Quantitative RT-PCR analysis revealed that NMDAR subunits are present in the aortic samples (Fig. 4 a, b). Hence, we investigated the glutamate receptor expression in isolated endothelial cells. Aortic endothelial cells (aEC) as well as brain endothelial cells (bEC) express NMDAR subunits with NR1 and NR2a in higher amounts compared to gliomas (Fig. 4 c, d). Consistent with previous reports, immune-cytochemical investigations revealed that CD31-positive endothelial cells derived from aorta and brain both express the NMDAR pan-subunit NR1 (15) (16) (Fig. 4 e). Moreover, we demonstrate that the NMDAR subunit NR1 is expressed in the vasculature of retinas and brain sections (Fig. 4 f).

**FIGURE 4:**
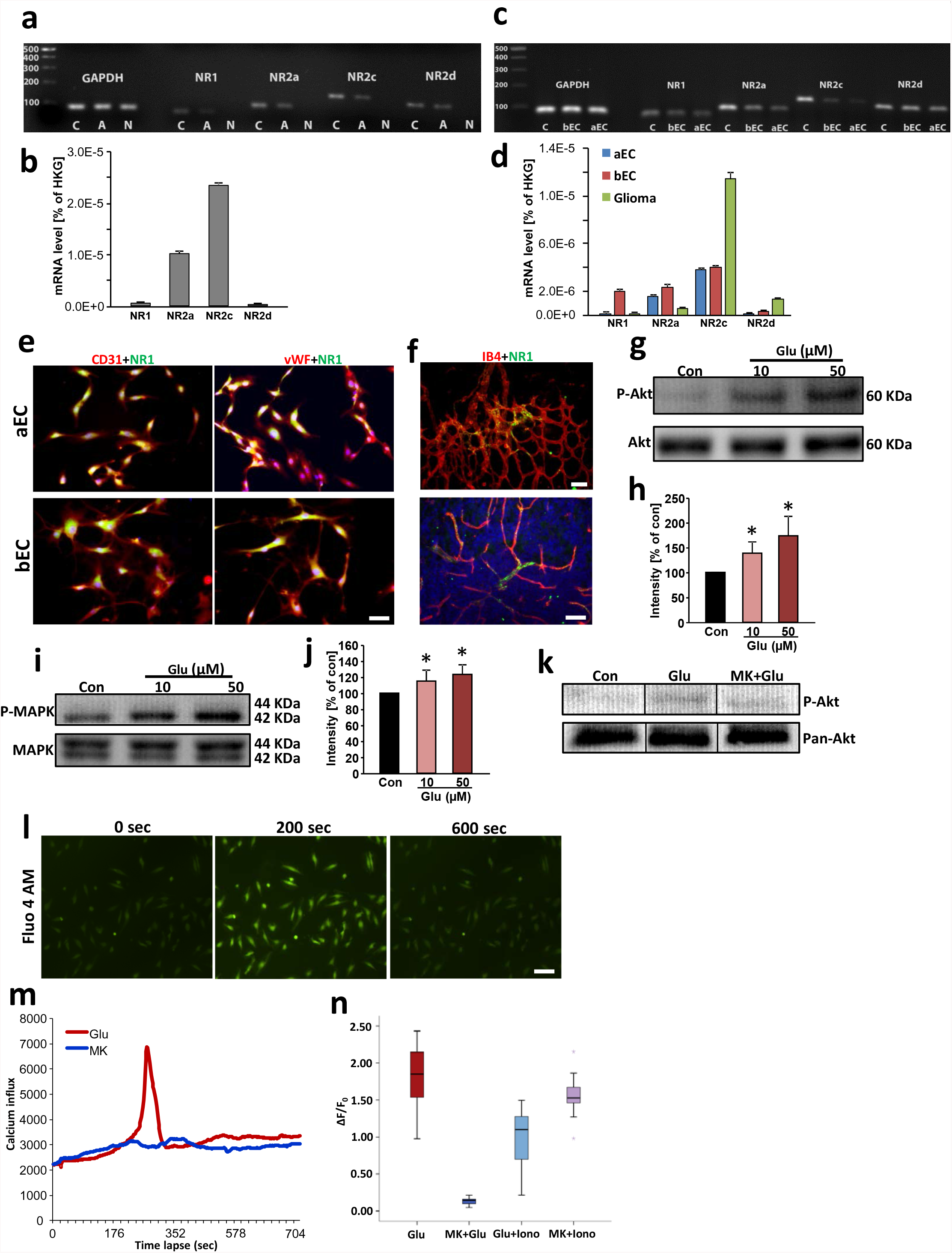
Glutamate induces angiogenesis via endothelial NMDA receptor activation. **a**, NMDA receptor subunit detection in rat endothelial cells. Agarose gel electrophoresis of RT-PCR amplicons of NR1, NR2a, NR2c and NR2d from rat cerebral cortex serving as a as positive control (C), aorta (A) and negative control (N). The housekeeping gene glyceraldehyde 3-phosphate dehydrogenase (GAPDH) was facilitated as an internal control. **b**, Quantitative real time RT-PCR for NMDA receptor subunit NR1, NR2a, NR2c and NR2d expression in rat aorta. The respective mRNA expression values are given as ratios to the house keeping gene (HKG) expression level GAPDH (means ± s.d.; n = 3 per group). **c**, NMDA receptor subunits are expressed in isolated rat brain endothelial (bEC) and rat aorta endothelial cells (aEC). Agarose gel electrophoresis of RT-PCR amplicons of NR1, NR2a, NR2c and NR2d are shown from rat brain serving as a positive control (C), brain endothelial cells (bEC) and aorta endothelial cells (aEC). The housekeeping gene glyceraldehyde 3-phosphate dehydrogenase (GAPDH) was facilitated as an internal control. **d**, Quantitative real time RT-PCR for NMDA receptor subunit NR1, NR2a, NR2c and NR2d expression in rat brain endothelial cells (BE), rat aorta endothelial cells (AE) and F98 glioma cells. The respective mRNA expression values are given as ratios to the house keeping gene expression level of GAPDH (means ± s.d.; n = 3 per group). **e**, Brain and aorta endothelial cells express the pan-NMDAR subunit NR1. Representative images of isolated aorta endothelial cells (aEC) (top raw) and rat brain endothelial cells (bEC) (bottom raw) immuno-stained for NR1 (green) and CD31 or von Willebrand factor (vWF) facilitated as endothelial cell marker (both shown in red). Nuclei were stained with Hoechst (blue). Scale bar represents 50 µm. **f**, Retina and brain vasculatures express the pan-NMDAR subunit NR1. Representative images of mouse retina (top) and mouse brain cryosections (bottom) immuno-stained for NR1 (green) and isolectin-B-4 facilitated as endothelial cell marker (shown in red). Nuclei were stained with Hoechst (blue). Scale bars in top represents 100 µm, in bottom 50 µm. **g**, Akt phosphorylation after glutamate stimulation in brain endothelial cells (RBEC). Serum-starved RBECs were untreated (con), or treated with various concentrations of glutamate (Glu 10 µM, 50 µM) for 30 minutes and subsequently, proteins were isolated for immunoblotting. Representative Western blot is shown demonstrating phospho-Akt dynamics in brain endothelial cells in response to glutamate normalized to total Akt levels (bottom). **h**, Quantification of p-Akt/total Akt levels expressed as the average percentage increase (mean ± s.d.) over basal levels (100%), Statistical significance was calculated with one way ANOVA, n = 3; *P < 0.05 versus control. **i**, MAPK phosphorylation after glutamate stimulation in brain endothelial cells (RBEC). Serum-starved RBECs were untreated (con), or treated with various concentrations of glutamate (Glu 10 µM, 50 µM) for 30 minutes and subsequently, proteins were isolated for immunoblotting. Representative Western blot is shown demonstrating phospho-MAPK dynamics in brain endothelial cells in response to glutamate normalized to total MAPK levels (bottom). **j**, Quantification of p-MAPK/total MAPK levels expressed as the average percentage increase. Statistical significance was calculated with one way ANOVA, n = 3; *P < 0.05 versus control. **k**, NMDA receptor antagonist MK801 blocks glutamate-induced Akt phosphorylation, Representative western blot is shown demonstrating phospho-Akt dynamics in brain endothelial cells in response to glutamate and MK801 normalized to Akt levels (bottom). **l**, Glutamate treatment triggers calcium transients in endothelial cells, most probably as influx through NMDAR. Representative images of HUVEC cells loaded with calcium indicator Fluo-4 AM. Green fluorescence indicates the level of intracellular calcium at different time points after glutamate treatment. **m**, Time lapse analysis of Ca^2+^ response in glutamate stimulated (Glu, 100 µM) HUVECs. MK801 application (100 µM) plus 100 µM glutamate (Glu + MK) served as controls in HUVECs. Note that calcium influx peak occurred at about 200 seconds after glutamate treatment, while MK801 fully blocked calcium influx. **n**, Quantification of ΔF/F_0_ shown in box plots. Calcium Ionophore A23187 was used as positive control to determine the highest level of Ca^2+^ influx by Ca^2+^-permeabilizing the plasma membrane.

These data imply that the excitatory neurotransmitter glutamate may generally impact on endothelial cell signaling. Therefore, we tested the downstream effectors activated by NMDAR signaling (17) (18). In response to different glutamate concentrations, endothelial cells showed increased levels of activated Akt (p-Akt) (Fig. 4 g, h). In addition, we tested whether NMDAR-dependent MAPK activation is inducible in endothelial cells. Therefore, we applied various concentrations of glutamate on endothelial cells and determined the NMDA downstream effector MAPK. We found that glutamate induced elevated phosphorylation of MAPK compared to controls, indicating a functional NMDA receptor activation (Fig. 4 i-j). To further test whether glutamate acts in a receptor-mediated manner on endothelial cells via NMDA activation, we pretreated endothelial cells with MK801 before glutamate application. These experiments showed that Akt activation was reduced following NMDAR inhibition, confirming that glutamate mediates receptor-dependent signals in endothelial cells (Fig. 4 k). We further investigated NMDAR-dependent responses in endothelial cells. Glutamate application on endothelial cells evoked a significant Ca^2+^ release (Fig. 4 l). Importantly, this glutamate-dependent Ca^2+^ response could be abolished by application of the NMDAR inhibitor MK801 (Fig. 4 l, m). Altogether, these data indicate that endothelial cells transduce glutamate signaling through NMDAR downstream kinases and Ca^2+^ responses.

### Inducible NMDAR1 knockout mice reveal compromised endothelial sprouts and vessel density

To study the role of glutamate-mediated angiogenesis *in vivo*, and to circumvent the embryonic lethality of a global knockout of the NMDA receptor subunit NR1 (GRIN1) (19), we generated inducible endothelial-cell specific genetic loss-of-function mutants (GRIN^iΔEC^). LoxP-flanked NMDAR1 allelic mice (GRIN^fl/fl^) were crossed with transgenic mice expressing the tamoxifen-inducible recombinase CreERT2 under the control of the endothelium-specific Cdh5 promoter. First, we tested the angiogenic potential of endothelia in aortic explant cultures. Following tamoxifen administration, controls displayed increased sprout formation and vascular density in the presence of glutamate (Fig. 5 a, b). In contrast, GRIN^iΔEC^ mutants showed significantly reduced endothelial sprouting and endothelial density (Fig. 5 a, b). Moreover, induced endothelial-cell-specific deletion of NMDAR1 between postnatal day 1 (P1) and P4 revealed that endothelial sprouting and vascular meshes were strongly reduced in GRIN^iΔEC^ retinas after glutamate treatment compared to control littermates (Fig. 5 c, d). Moreover, endothelial-cell-specific deletion of NMDAR1 significantly reduced the number of endothelial filopodia *in vivo* (Fig. 5 e, f).

**FIGURE 5:**
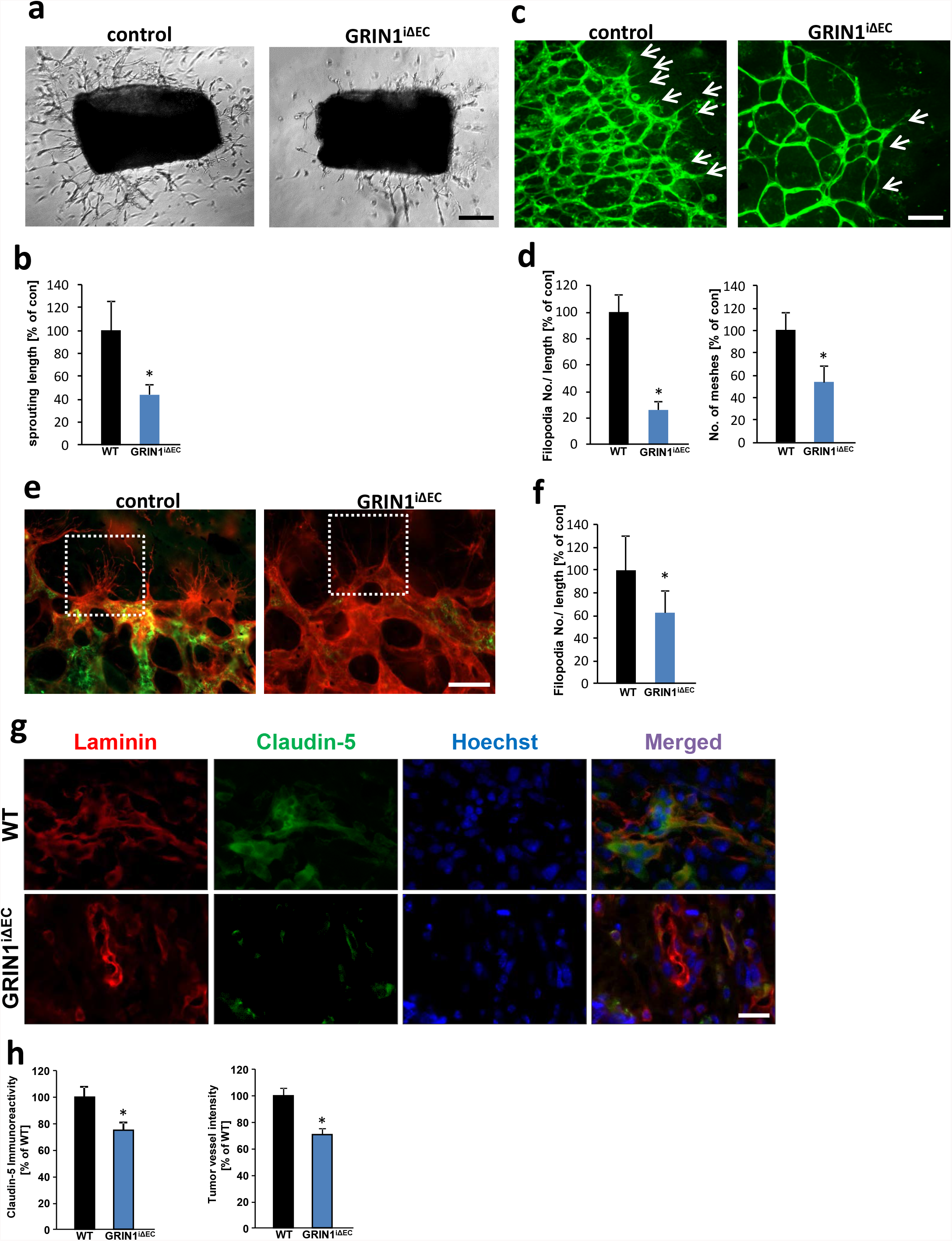
Endothelial-specific NMDAR inhibition diminishes angiogenesis. **a**, Aortic explant cultures from control mice and endothelial-cell-specific NMDAR1 mutants (GRIN^iΔEC^) in the presence of 100 µM glutamate. Representative images of aortic explants show endothelial sprouts which are reduced in GRIN^iΔEC^ mutants compared to controls. Scale bar represents 200 µm. **b**, Quantification of endothelial sprouting defects in GRIN^iΔEC^ mutants compared to controls. Statistical significance was calculated with student’s t. test, (means ± s.d., *P < 0.05, from n = 12). **c**, Isolectin-B4 stained control and GRIN^iΔEC^ retinas (tamoxifen treated from P1 to P4) in the presence of 100 µM glutamate. Arrows indicate endothelial sprouts. Scale bar: 100 µm. **d**, Quantification of vascular parameters of retinas (filopodia numbers/ retina length; endothelial meshes), control group was set to 100%. (n ≥ 3). **e**, Isolectin-B4 (red) and NR1 antibody (green) stained control and GRIN^iΔEC^ retinas (tamoxifen treated between P1 and P7). White dashed box focuses endothelial filopodia. Scale bar: 50 µm. **f**, Quantification of retinas filopodia numbers/ retina length, control group was set to 100%. (n ≥ 3). **g**, Representative immunofluorescence images for Laminin (red) and Claudin-5 (green) in the peritumor area of gliomas implanted in wild type control mice (WT) and endothelial-cell-specific NMDAR1 knock out mice (GRIN^iΔEC^). Nuclei are stained with Hoechst and are given in blue. Scale bar represents 100 µm. **h**, Quantification of peritumor Claudin-5 immunoreactivity and vessel density in wild type mice (WT) and endothelial-cell-specific NMDAR1 knock out mice (GRIN^iΔEC^). Values of wild type mice group is given as 100% (n = 3 per group).

Hence, we investigated whether NMDAR expression in brain endothelial cells has an impact on tumor-induced angiogenesis. Therefore, we implanted syngeneic gliomas orthotopically into brains of wildtype controls and GRIN^iΔEC^ mice and monitored tumor vessels (Fig. 5 g). Gliomas in NMDAR knockout background (GRIN^iΔEC^ mice) displayed significantly reduced tumor vessels compared to controls (Fig. 5 g, h). Altogether, these data show that NMDAR signaling is regulating endothelial sprouting and affect tumor-induced angiogenesis.

### xCT inhibition in brain tumors normalizes tumor angiogenesis

Since the expression of xCT correlates with shorter glioma patient survival (7) (NCB molecular brain neoplasia database, data not shown) and the malignancy of gliomas correlates with the degree of angiogenesis, we studied the tumor effects of xCT *in vivo*. We compared the net tumor volume effects in xCT overexpressing and knockdown gliomas (Suppl. Fig. 4). By MRi scans we determined the tumor-induced brain edema and found that xCT^KD^ gliomas showed alleviated brain edema compared to WT gliomas (Suppl. Fig. 4). In contrast, xCT^OE^ gliomas displayed increased brain edema and thereby increased space-occupying lesions (Suppl. Fig. 4).

We further investigated whether therapeutic xCT inhibition could contribute to tumor vessel normalization (vessel parameters equal to controls) *in vivo*. For this, we implanted syngeneic gliomas orthotopically into brains of animals and monitored vessel architecture by synchrotron X-ray tomographic microscopy (20) (21). Wild-type gliomas displayed pathological vascular alterations in the peritumor and tumor zones in comparison to the contralateral hemisphere without tumor affection serving as a control (Fig. 6). In particular, vessel branches and lengths of metarterioles and capillaries were significantly altered within the tumor bulk as well as in peritumor regions (Fig. 6 a-c). In contrast, siRNA-mediated knock down of xCT in gliomas normalized the vascular pattern (vessel branches and length) resembling those of controls (Fig. 6 c-e, purple underlined area). Metarterioles and capillaries resembled a phenotype closely to that found in non-affected control brain regions (purple boxed area) (Fig. 6 c-e). These findings demonstrate that blocking glutamate secretion abrogates glutamate-triggered tumor vessel growth.

**FIGURE 6:**
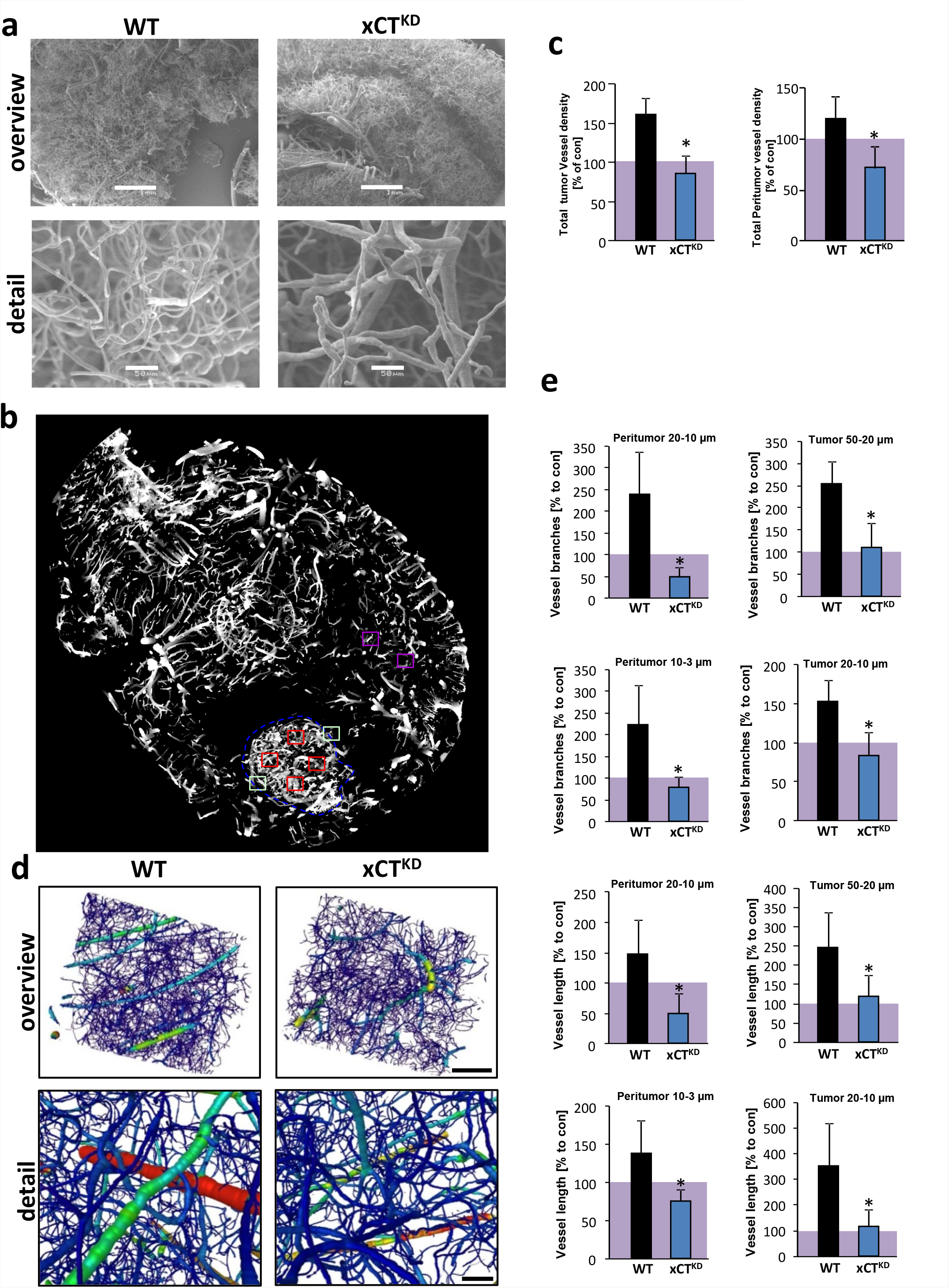
RNAi-mediated xCT silencing in glioma cells alleviates tumor-induced angiogenesis in vivo. **a**, Representative images from the scanning electron microscopy of brain vessel casts with tumor-implantation of wild-type gliomas (WT) and xCT^KD^ gliomas (xCT^KD^). Scale bars: 1 mm in overview images and 50 µm detail images. **b**, Representative 2-D image of synchrotron X-ray microscopic tomographic slice of a tumor-implanted brain specimen. Positions of quantitation of vessels in the tumor bulk (red squares), peritumor zones (green squares) and control areas (violet). **c**, Quantitative analysis of overall vessel density in tumor and peritumor regions. Purple area represents values from contralateral brain areas without tumor affection (n ≥ 5 per group). **d**, Representative images of a reconstruction of synchrotron X-ray microscopic tomographic slices of brains with wild-type gliomas (WT) and xCT^KD^ gliomas. Blood vessel objects were visualized and color coded according to the vessel diameter from red (thick) to purple (thin). **e**, *Left* column, Quantitative analysis of the vascular architecture in the peritumor zone of wild-type (WT, black column) and xCT knock down (xCT^KD^, blue column) gliomas. Vessel branch numbers and vessel lengths were analyzed for metarterioles (diameter Ø 10–20 µm) and capillaries (Ø 3–10 µm) compared to control vascular brain regions (control values are highlighted in purple underlined area). Branch number and length of metarterioles (Ø 10–20 µm) and capillaries (Ø 3–10 µm) are significantly decreased in xCT^KD^ gliomas. *Right column*, Analysis of the intratumor vascular architecture of wild-type (WT, black column) and xCT knock down (xCT^KD^, blue column) gliomas. Vessel branch numbers and vessel lengths were analyzed and arterioles (Ø 20–50 µm) and metarterioles (Ø 10–20 µm) quantified in comparison to control vascular brain regions. WT glioma values are given in black columns, xCT^KD^ glioma values are given in blue columns and control values are highlighted in purple. Statistical significance was calculated with student’s t. test, (means ± s.d., *P < 0.05, from n ≥ 5).

## DISCUSSION

The expression level of the glutamate-cysteine antiporter xCT has an important role in determining the malignancy in different human tumors entities (7) (6) (22) (12, 14). Here, we investigated the underlying mechanisms of xCT in malignant gliomas. We provide evidence for a functional role of the glutamate and NMDAR signaling pathway in tumor vessel formation.

Since tumor-derived glutamate release is one major factor contributing to tumor-associated neurodegeneration, xCT modelling represents an excellent prerequisite to study the causal relation between neurotoxic events and vascular responses. Fostered xCT expression increased peritumoral vascularization and the development of irregular vessel structures. Similar effects were found by applying the neurotransmitter glutamate into the brain microenvironment. Thus, our data indicate that the effects of xCT on vessels are mediated through discharged glutamate and potential glutamate receptor-dependent signaling in endothelial cells. In fact, the biological effects of glutamate were shown to be in accordance with clinical measurements of peritumoral glutamate in glioma patients, which reached excess levels of up to 100 µM in human patients (10). Importantly, the increase in excitatory neurotransmitter glutamate levels is toxic for neurons (23) (9), whereas endothelial cells are resistant to high glutamate concentrations and endothelial viability remained unaffected (15). In addition, our data and previous reports show that the difference between endothelium and neurons is not due to lack of glutamate signaling in brain vessels (24). Independent lines of evidence already demonstrated that endothelial cells express functional NMDARs and showed that glutamate is involved in regulating arteriole diameter, vasodilation and blood brain barrier disruption (15) (25) (16, 26). Our data indicate that xCT-dependent effects are mediated by NMDAR signaling and are reversible by the action of NMDAR antagonists. These findings are further supported by inducible loss-of-function genetics in endothelial cells *in vivo*. We generated an inducible endothelial cell-specific NR1 knockout model due to the circumstances that conventional NR1 gene disruption causes neonatal lethality (27, 28). Endothelial cell-specific NR1 knockout revealed that glutamate receptor signaling contributes to endothelial sprouts. This strengthens the evidence for the involvement of NMDA receptors mediating pro-angiogenic effects of glutamate by pharmacological and loss-of-function genetic experiments. Thus, xCT in gliomas contributes not only to cytotoxic events but also to vasogenic processes which both converge into tumor-associated brain swelling. This study further supports a previously discovered neuro-vascular crosstalk of the neurotransmitter glutamate and vascular signaling. In fact, it has been shown that the vascular growth factor receptor 2 is expressed in neurons and forms a complex with NMDAR subunits mediating neuronal migration and survival (29) (30) (31) (32). Brain tumors may take advantage of this signaling convergence by facilitating xCT which operates on neuronal and endothelial integrity. Our results demonstrated that glutamate has a direct pro-angiogenic impact on endothelial cells via NMDAR signaling. Finally, in a therapeutic intervention, we analyzed the vascular response to reduced glutamate through xCT inhibition. Malignant brain tumors with silenced xCT expression showed a normalized tumor vasculature and reduced aberrant vessel formation supporting glutamate and NMDAR as a target for anti-cancer treatment. This is particularly important since sole xCT inhibition can neither pharmacologically nor genetically reduce glioma growth significantlly, although it alleviates clinical malignancy characteristics (Robert et al., 2015; (33); (6, 9).

In conclusion, our data demonstrate that the glutamate transporter xCT and the glioma-derived neurotransmitter glutamate have a crucial role in tumor angiogenesis. Thereby, glutamate receptors of the NMDA subclass operate in endothelial cells and convert the neurotransmitter glutamate to a promoter of angiogenesis. Hence, these data give unique insights into the molecular basis of xCT-mediated tumor angiogenesis and provide fundamental understanding for the biology of glutamate in vascular signaling.

## MATERIALS AND METHODS

### Chemical, drugs and cell lines

Rodent glioma cell lines F98, C6, and the human glioma cell line U251 were obtained from ATCC/LGC-2397 (Germany) and were cultured under standard condition containing DMEM medium (Biochrom, Berlin, Germany) supplemented with 10% fetale bovine serum (FBS) (Biochrom, Berlin, Germany), 1% Penicillin/Streptomycin (Biochrom, Germany) and 1% Glutamax (Gibco/Invitrogen, California, USA). Cells were passaged at approx. 80% confluence by adding trypsin after 1 PBS wash step and incubated for 5 min, then centrifuged at 900 rpm/5 min. Cell lines were transfected as described previously (Savaskan et al., 2008) and maintained under standard conditions. GYKI 47261 dihydrochloride (4-(8-Chloro-2-methyl-11H-imidazo[1,2-c][2,3]benzodiazepin-6-benzeneamine dihydrochloride) and MK801(Dizocilpine) were purchased from Tocris Biosciences (R&D Systems GmbH, Wiesbaden, Germany).

### RNA isolation and quantitative RT-PCR analysis

Cells were collected when grown at subconfluency, followed by suspension in phosphate buffered saline (PBS). Total RNA was extracted using High Pure RNA Isolation Kit (Roche, Mannheim, Germany) following the manufactory’s manual. RNA concentration was determined by NanoVue™ Plus Spectrophotometer (GE Healthcare, UK). cDNA synthesis was performed with SuperScript® III Reverse Transcriptase (Invitrogen). qRT-PCR was performed with SYBR Green PCR master mix (Qiagen). The oligonucleotides used in this study are: xCT forward primer: TGCTGGCTTTTGTTCGAGTCT; xCT reverse primer: GCAGTAGCTCCAGGGCGTA. GAPDH forward primer: TGCACCACCAACTGCTTAGC; GAPDH reverse primer: GGCATGGACTGTGGTCATGA. NR1 (Cat.-No.: QT01751484), NR2a (QT01588069), NR2c (QT01751498), NR2d (QT00193438), VEGFa (QT00198954), VEGFb (QT01290163) primers were purchased from Qiagen (Germany), Beta-actin forward primer: GCTCCTCCTGAGCGCAAG; Beta-actin reverse primer: CATCTGCTGGAAGGTGGACA. Real time cycling parameters were: Initial activation step (95°C, 15 min), cycling step (denaturation 94°C, 15 s; annealing at 60°C, 30 s; and finally extension for 72°C, 30 s X40 cycles), followed by a melting curve analysis to confirm specificity of the PCR. The Ct value was corrected by Ct reading of corresponding GAPDH or β-actin controls. Data from three determinations (means ± SEM) are expressed as relative expression level. The reaction was performed using Light Cycler 480 (Roche). The specificity of the PCR reaction was confirmed by agarose gel electrophoresis.

### Protein isolation and immunoblotting

For protein analysis cells were seeded in 6-well plates. Cells were maintained under standard conditions for 3 days afterwards medium was removed and cells washed with PBS for one time. Then, 1 ml supplements-free ECGM medium was added and cells incubated in 37°C for 30 minutes. Glutamate treatment was conducted by adding 10 µl out of a glutamate stock of 1 mM and 5 mM into the supplements-free ECGM. After 30 minutes incubation, medium was sucked off and quickly lysed with NP 40 buffer containing 5 mM NaF and a protease inhibitor cocktail (Roche, Basel, Switzerland) and homogenized by ultrasound (Bandelin Sonoplus, at 67%). After 20 min on ice, samples were centrifuged at 8000 rpm for 8 min. Supernatants were isolated and protein content measured with the NanoVue™ Plus Spectrophotometer (GE Healthcare, UK). Samples were mixed with loading buffer (4x) and reducing agent (10x) (Invitrogen, California, USA) and boiled at 96°C for 8 min. Equal amounts of protein sample were loaded on 4-12% SDS-NuPage gel (Invitrogen, CA, USA) and electrophoresis was performed in MOPS-buffer, transferred on PVDF membranes (Roth, Karlsruhe, Germany). Transfer efficiency and consistency was checked with Memcode Stain kit (Thermo Massachusetts, USA) according to the user manual. Membranes were blocked in PBS containing 2% Magic block and 10% Top block (Lubio science, Luzern, Switzerland) for 1h before further processed. Antibodies were incubated overnight at 4°C in roller tubes, followed by secondary antibodies incubated at room temperature for 1 hour. Detection was performed with ECL plus kit (GE-healthcare, Solingen, Germany).

### Expression vectors and knockdown vector cloning

Reverse transcription-polymerase chain reaction was used for full length cloning of xCT from rat, mouse and human mRNA samples (9). For sequence alignments and homology searches of xCT we utilized the www.ncbi.nlm.nih.gov database and A Plasmid editor software (ApE; MW Davis, Utah, USA). All orthologous sequences of xCT (human, mouse and rat) are deposited at the NCBI database (Human xCT GenBank accession no. AF252872; Rattus norvegicus xCT GenBank accession no. NM001107673; Mus musculus xCT GenBank Accession no. AB022345). For construct cloning we cloned fragments by PCR and inserted the resulting amplicons into the pEGFP (Takara, Heidelberg, Germany) vector. According to the critera of Ui-Tei et al. (34) three 19-mer short interfering RNAs were chosen for RNA interference with rat xCT transcripts (GenBank acc. NM001107673). Cloning of the synthetic oligonucleotids into the pSuperGFP vector (pS-GFP; OligoEngine) was performed by digesting the empty vector with EcoR1 and Xho1 according to the manufacturer’s instruction. Cells were transfected at low density (<20.000 cells/cm²) and expression analysis was performed as described (6).

### Mutant mice and inducible genetic deletion experiments

For endothelial-cell specific genetic loss-of-function experiments, we inbred *loxP*-flanked *Grin1* (*Grin1*^lox/lox^) mice (35) with transgenic mice expressing the tamoxifen-inducible recombinase CreERT2 under the control of the endothelial Cdh5 promoter (*Cdh5 (PAC)-CreERT2*) (36). Cre activation was induced by tamoxifen injection as previously described (37). Briefly, for adult mice, intraperitoneal injection of 100 µl of a 20mg/ml tamoxifen solution was applied daily for a duration of 4 days. For new born mice, 50 µl tamoxifen (1mg/ml) was injected into mice stomach once daily from postnatal 1 to postnatal 4.

### Endothelial cell isolation

For isolating endothelial cells from rodents and mice, brain tissue and thoracic aorta were dissected and transferred into dishes with sterilized PBS buffer. Tissue sections were placed in ECGM and cultured on the Matrigel-coated wells until microvessel-like out-sprouts migrated. The brain endothelial and aortic endothelial cell cultured revealed in 95% positive for CD31 (PECAM-1) and Von Willebrand factor (vWF). Human umbilical vein endothelial cells (HUVECs) were isolated from freshly collected umbilical cords by standard techniques as described (38).

### Tube forming assay and aortic ring explant cultures

96-well plates were evenly coated with 30 μl/well 60% matrigel (BD Biosciences) (60% matrigel + 40% DMEM) and incubated at 37 °C for 1 hour until the matrigel of each well became polymerized. Endothelial cells were seeded in a volume of 100 μl endothelial growth medium for each well and treatment was performed 30 minutes afterwards. The total length of tubes was quantified by angiogenesis plug in tools of Image J (NIH). For aortic ring assay, 1 month old Wistar rats were sacrificed by head dissection. Thoracic aortas were dissected and transferred into a dish with sterilized PBS buffer. After removing the fibroadipose tissue, aortic rings were cut and placed on collagen-coated 96-well plates and embedded in matrigel. Endothelial sprouting length was measured in an automatized fashion using NIH-Image J software.

### Retinal explant cultures

For retinal cultures, we followed the protocol according to Sawamiphak et al., 2010 and Pitulescu et al., 2010 (39). Briefly, postnatal day 4 mice were sacrificed by quick head dissection and the retina was isolated. Retinas were then transferred onto culture plate insert membrane dishes of six-well culture dishes (GreinerBioOne) containing 1.2 ml culture medium (DMEM, Biochrom). After 4 hours of treatment in 35°C, 5% CO_2_, the retina was fixed by 4% PFA for 30 min at room temperature and stained with isolectin GS-IB4 (1:250) (Life technologies, Germany)

### Vascular organotypic glioma invasion (VOGIM) brain cultures

Brain slice cultures were conducted with five days old Wistar rats. Brains were prepared and maintained as previously described (40). Briefly, animals were sacrificed by quick head dissection and brains were removed and kept under ice-cold conditions. Frontal lobes and Cerebellum were dissected of the hemispheres. The remaining brain was cut into 350 µm thick horizontal slices using a vibratome (Leica VT 1000S, Bensheim, Germany). The brain slices were thereafter transferred onto culture plate insert membrane dishes with a pore size of 0.4 µm (GreinerBioOne, Frickenhausen, Germany) and subsequently transferred into six-well culture dishes (GreinerBioOne) containing 1.2 ml culture medium (MEM–HBSS, 2:1, 25% normal horse serum, 2% L-glutamine, 2.64 mg/ml glucose, 100 U/ml penicillin, 0.1 mg/ml streptomycin, 10 µg/ml insulin–transferrin–sodium selenite supplement and 0.8 µg/ml vitamin C). Slices were cultured in humidified atmosphere (35°C, 5% CO_2_). After 24 hours, slices were gently washed with 1.2 ml PBS and a full culture medium exchange was performed. At the second day after preparation, 0.1 µl of the cell-medium-suspension (containing 10,000 cells) was placed onto the temporal cortex of the slice by using a 1-µl-Hamilton-syringe. The medium was exchanged every second day. For MK801 treatment, 1.2 µl 1 mg/ml MK801 was applied to the medium reaching final concentration of 100 µM MK801. Slices were fixed and stained for laminin (Sigma) (1:250) for vessel analysis.

### Organotypic vascular sponge assay

Agarose cubes were made from 1% agarose containing only PBS (control), 100 µM glutamate or 200 µM glutamate. Cooled agarose hydrogels were then cut into approximately 1mm cubes and implanted on the cortex area of rat brain slices as described. Cube-implanted brain slices were cultured on semipermeable membrane dishes with a pore size of 0.4 µm (GreinerBioOne, Frickenhausen, Germany) and subsequently transferred into six-well culture dishes (GreinerBioOne) containing 1.2 ml culture medium. The slices were cultured in humidified atmosphere (35°C, 5% CO_2_). Media change was performed every other day. Four days after sponge cube implantation, slices were fixed and stained with anti-Laminin-antibody (Sigma) (1:250) for vessel analysis.

### Intravital cerebral tumor-vessel microscopy

Animal experiments were performed according to local animal welfare guidelines. Adult athymic nude mice (Jax: 002019) were anesthetized deeply. Anesthesia was verified by foot paw pinching reflex control. Mouse cranial window by craniotomy was described previously (41). Briefly, the mice were stereotactically fixated and a 6 mm Ø craniotomy was performed using a micro-drill (0.5 mm, Figure 3c). The Dura matter was carefully removed using micro forceps, the tumor spheroid was carefully placed on a vessel free area of the cortex. The window was sealed with a 7 mm Ø cover glass and the skin was sutured. Intravital epifluorescence video microscopy was performed 7, 10, and 14 days following implantation using a modified Axiotech vario microscope (Attoarc; Zeiss, Germany) with 10× long-distance and 20× water-immersion objectives (Zeiss) 40. DiI labeled glioma cells grown as sphenoids for precise delineation of the tumour from the adjacent host tissue by epifluorescence (520–570 nm) (Laib et al., 2009). Microvessels were visualized by contrast enhancement with 2% FITC-conjugated dextran (0.1 mL, intravenous; molecular weight 150,000; Sigma) and the use of blue-light epi-illumination (450–490 nm). Microscopic images were recorded through a charge-coupled device (CCD) video camera with an optional image intensifier for weak fluorescence (Kappa) and transferred to a S-VHS video system (Panasonic, Japan) for offline analysis. Offline analysis was performed using a computer-assisted analysis system (CAPIMAGE; Zeintl Software Engineering, Heidelberg, Germany). To assess vascular permeability we compared intravascular to extravascular fluorescence intensity of multiple individual host and tumour blood vessels and calculated a permeability index. For microcirculatory analysis, the newly formed microvasculature of the tumour was assessed by 4-5 ROI measurements per animal and day. Quantitative analysis included functional intratumoural vascular density (FVD), defined as the length of red blood cell perfused microvessels, vessel diameter (D), microvascular permeability (P), calculated as the ratio between intra and extravascular contrast and reflecting the extent of fluorescent marker extravasation, and microvascular red blood cell velocity (RBCV; mm/s).

### In vivo tumor cell implantation

Fischer 344 rat were stereo-tactically injected with F98 rat glioma cells stably expressing scrambled shRNA or pEGFP vector (control), pEGFP-xCT and xCT specific shRNA (xCT^KD^) constructs as described previously (6).

### Frozen sections and immunofluorescence

Rat brains were isolated 11 days after tumor implantation and fixed in 4% PFA overnight at 4 °C. Thereafter, fixed brains were washed with PBS and incubated in 20% sucrose for 3 days. The brains were frozen in liquid nitrogen and embedded in Tissue Tek. For vessel analysis cryosections (20 µm) were stained with Alexa Fluor 568 isolectin GS-IB4 at 1:500 (Invitrogen, Life Technologies). Detection of neurons was accomplished with anti-NeuN antibody (MAB377, Chemicon, Millipore) (1:100) and Alexa Fluor 568 goat anti-mouse IgG secondary antibody (1:1000) (Molecular Probes, Life Technologies, Darmstadt, Germany). Cryosections were counterstained with HOECHST 33258 (Invitrogen, Life Technologies) at a concentration of 1 µg/ml.

### Scanning electron microscopy

Brain vessel corrosion casts from tumor-implanted rats were mounted on aluminum stabs and sputtered with 10 nm gold. The vasculature of the specimens was scanned with a Hitachi S4000 scanning electron microscope and subsequently analyzed with Image J (Heinzer et al., 2008).

### Synchrotron microscopic CT and vessel analysis

Eighteen days post implantation, rats were deeply anesthetized, killed and perfused with the PU4ii polymer resin (Vasqutec, Zurich, Switzerland) as described previously (42). Resulting vascular reconstructions of regions of interest (ROIs) of the contralateral hemisphere (control) peritumor and tumor cubes (740.16µm^3^ per ROI) were reconstructed using Amira Software (Visage Imaging, Andover USA). The cubes were cut into cylinders to prevent artifacts. Contrast dependent object identification was used to identify the blood vessels. Resulting data spreadsheets and self-designed programs and macros were processed using Microsoft Excel 2007. Vessels were categorized by diameter according to the anatomical classification of blood vessels: Arterioles (21–50 µm), metarterioles (11–20 µm) and capillaries (3 - 10 µm).

### Calcium imaging

Fluo-4 AM (Stock: 1mM in DMSO) was used in a final concentration of 2 µM. Approx. 10,000 HUVEC cells were seeded in 96 well plates and cultured overnight with 100µl/well in ECGM full medium (PELO Biotech). Endothelial cells were loaded with Fluo-4 AM at 37 °C for 30 min. Images were constantly recorded with Nikon TiE epifluorescence miscroscope for 15 min and calcium ionophore A23187 served as positive control (final concentration 10µM).

### Statistical analysis

Data from experiments were obtained from at least three independent experiments if not otherwise stated. Statistical analysis was performed using the ANOVA test using Graphpad Prism (GraphPad Inc., CA, USA) if not otherwise described in the figure legends. The level of significance was set at P∗ < 0.05, P ∗∗ < 0.01, P ∗∗∗ <0.001 according to the international conventions. Error bars represent ± S.D.

## NOTES

### Authors’ contribution

NES conceived and supervised the study. ZF performed the *in vitro* experiments with help of TB, TS and IYE. NES, TB and IYE performed the *in vivo* experiments with help from ZF, EPM and MC. *In vivo* experimentations with transgenic animals were conducted by ZF, EY, T.S:, NDS, MHH.S. and RN. Live cell imaging was planned and conducted by ZF, SS, and OF. TB and NES performed the Synchronton-XRTM analysis with help from MS. Data analysis and evaluation was conducted by ZF, TB, TS, SS, OF, IYE, NES and critically discussed with all authors. NES wrote the manuscript in conjunction with ZF, TB and IYE. All authors contributed to the preparation of the final manuscript and lent shape to the final paper.

## ACKNOWLEDGMENTS

We thank all members of the Translational Cell Biology & Neurooncology lab and in particular Ali Ghoochani, Nirjhar Hore and Daishi Chen (Erlangen, Germany) for valuable comments on this study and helpful suggestions. We are grateful to Ralph H. Adams (MPI Münster, Germany) for providing the Cdh5-Cre mouse line and Rolf Sprengel (MPI Heidelberg, Germany) for providing the GRIN^fl/fl^ mouse line. We thank Jan Lewerenz (Ulm, Germany) for various expression constructs, Peter Kalivas (Charleston, South Carolina, USA) for antibody exchange, Jan Csupor (Berlin, Germany), Christoph Hintermüller (ETH Zurich, Switzerland), Matthias Zenkel, and Ulrike Schlötzer-Schrehardt (both from Dept. of Ophthalmology, Erlangen, Germany) for excellent apparatus and technical support. We appreciate the prompt help with endothelial cell culture of A. Acker-Palmer (Frankfurt, Germany), and Doris Flick, Barbara Dietel and Matthias Beckmann (Erlangen, Germany) for providing initially HUVECs and technical support. Christiane Mühle and Martin Reichel (Department of Psychiatry, Erlangen, Germany) are gratefully acknowledged for providing assistance with the digital qRT-PCR procedure. We thank Manfred Rauh (Erlangen, Germany) for the mass spectrometric amino acid analysis and Hairuo Dang from D-HEST ETH Zurich for English editing and helpful suggestions. We are grateful to Robert Brandt (Visage Imaging, Berlin) for technical support on the Amira software and to the valuable work of Stefanie Gross and Gudrun Schöneberg (Dept. Dermatology, Erlangen, Germany) on the FACS application. Z.F. is supported by a grant from the China Scholarship Council (CSC: 2011627126). This study was funded by intramural (LOM) funding of the Universitätsklinikum and support of the, Verein zur Förderung des Tumorzentrums der Universität Erlangen-Nürnberg e.V to NES.

---

**SUPPLEMENTAL FIGURE 1:**
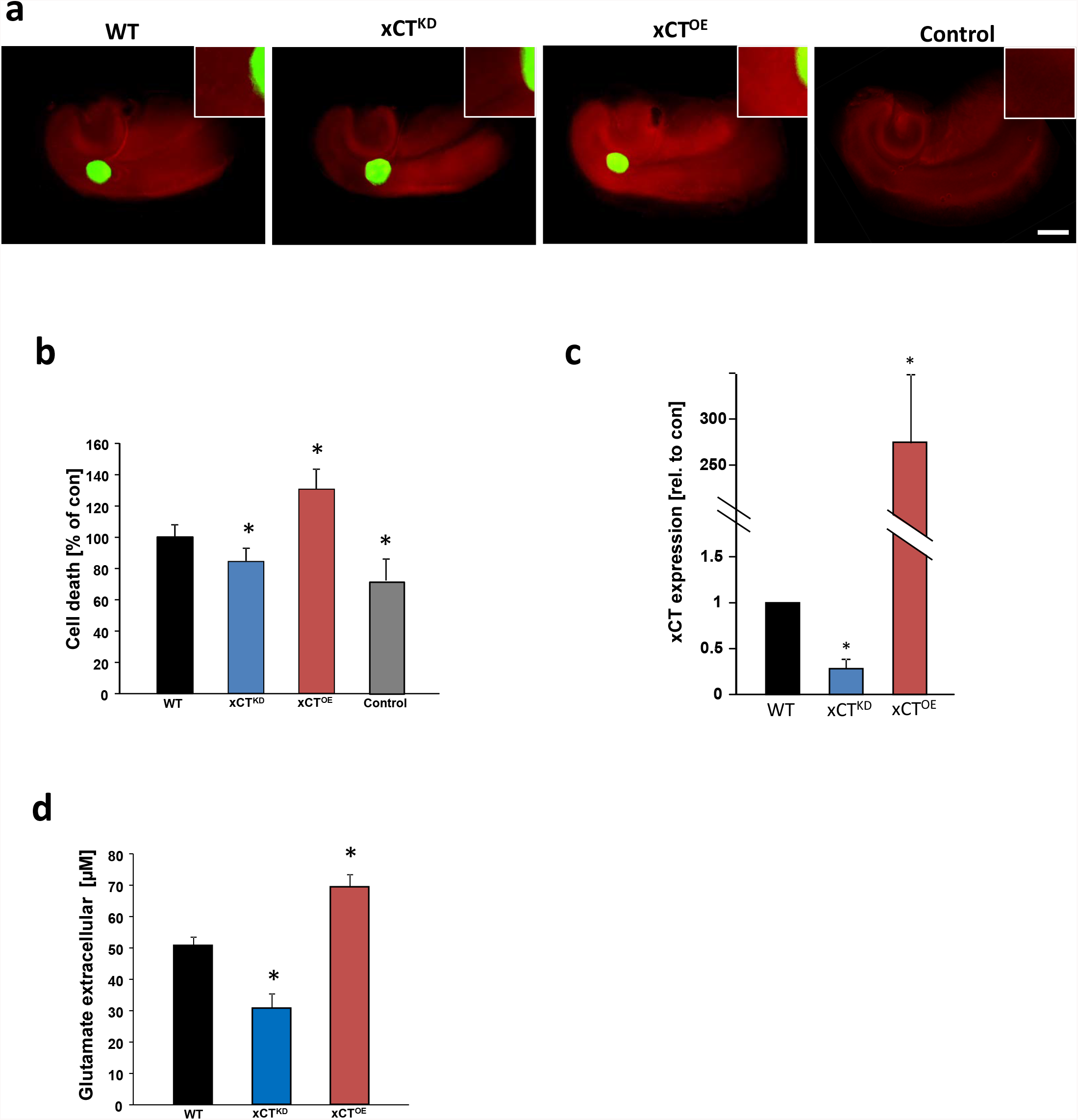
xCT-driven glutamate secretion induces neurodegeneration. **a**, RNAi-mediated xCT silencing alleviates tumor-induced neuronal cell death and xCT overexpressing tumors showed enhanced peritumour neurodegeneration. GFP^+^ glioma cells (scrambled siRNA expressing wild type gliomas, WT), xCT-knockdown glioma cells (xCT^KD^) and xCT overexpressing glioma cells (xCT^OE^) were implanted in brain tissue, and after 5 days cell death was evaluated (propidium iodide+ cells, red). Boxes on the top-right corner of each image display the magnification of peritumour area. Scale bar: 1 mm. **b,** Quantification of cell death intensity (propidium iodide+ signal). (n ≥ 6 per group). **c**, Quantitative real time PCR for xCT in scrambled siRNA expressing wild type gliomas (WT), xCT-knockdown glioma cells (xCT^KD^) and xCT overexpressing glioma cells (xCT^OE^). The respective mRNA expression values are given as ratios to the house keeping gene HPRT expression level, WT xCT expression level was normalized to 1. **d,** Extracellular glutamate levels were determined in glioma cells. HPLC results showed that medium from wild type (WT) glioma cells has 50.7 ± 2.8 µM glutamate. xCT silenced gliomas (xCT^KD^) showed decreased glutamate secretion (30.5 ± 4.5 µM) and xCT overexpressing gliomas (xCT^OE^) secrete elevated glutamate levels into the medium (69.4 ± 4 µM). (n = 3). *P < 0.05 versus control).

**Supplementary Figure 2:**
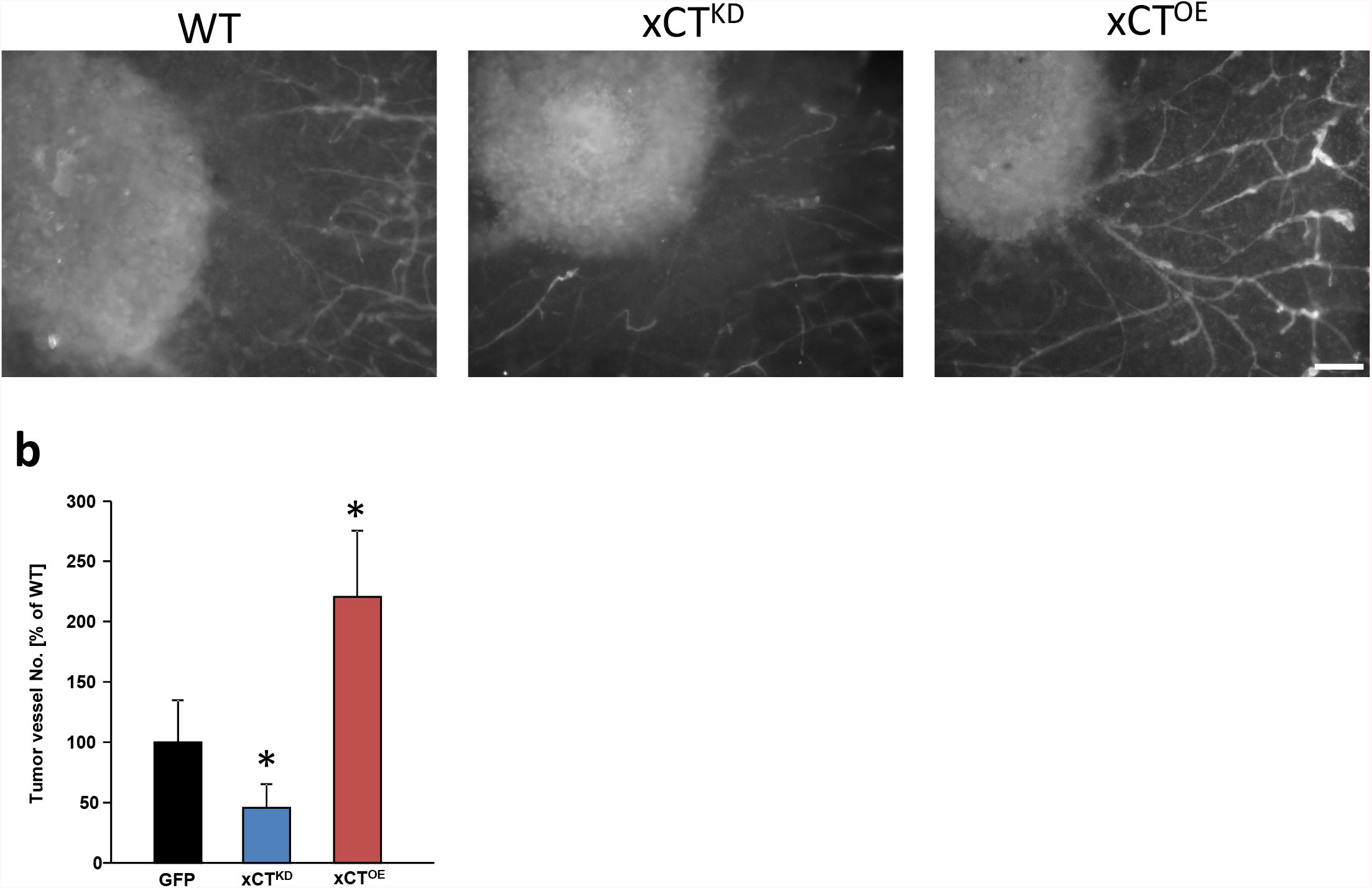
RNA-mediated xCT knockdown normalizes tumor angiogenesis in human gliomas. **a**, RNAi-mediated xCT silencing (xCT^KD^) alleviates human (U251) glioma-induced angiogenesis and xCT overexpressing U251 gliomas (xCT^OE^) show increased vessels. Representative images from scrambled siRNA expressing GFP^+^ gliomas (WT), xCT siRNA knock down (xCT^KD^) and xCT overexpressing (xCT^OE^) gliomas 7 days after tumor implantation. Vessels are immunostained for laminin. Scale bar represents 200 µm. **b**, Quantification of tumor vessels in peritumoral areas of WT, xCT^KD^ and xCT^OE^ U251 gliomas. Analysis was performed by counting the number of vessels which grow into the tumor. Control group was set as 100%. Statistical significance was calculated with one way ANOVA (means are given as percentage of controls ± s.d., *P < 0.05, n ≥ 3).

**SUPPLEMENTAL FIGURE 3:**
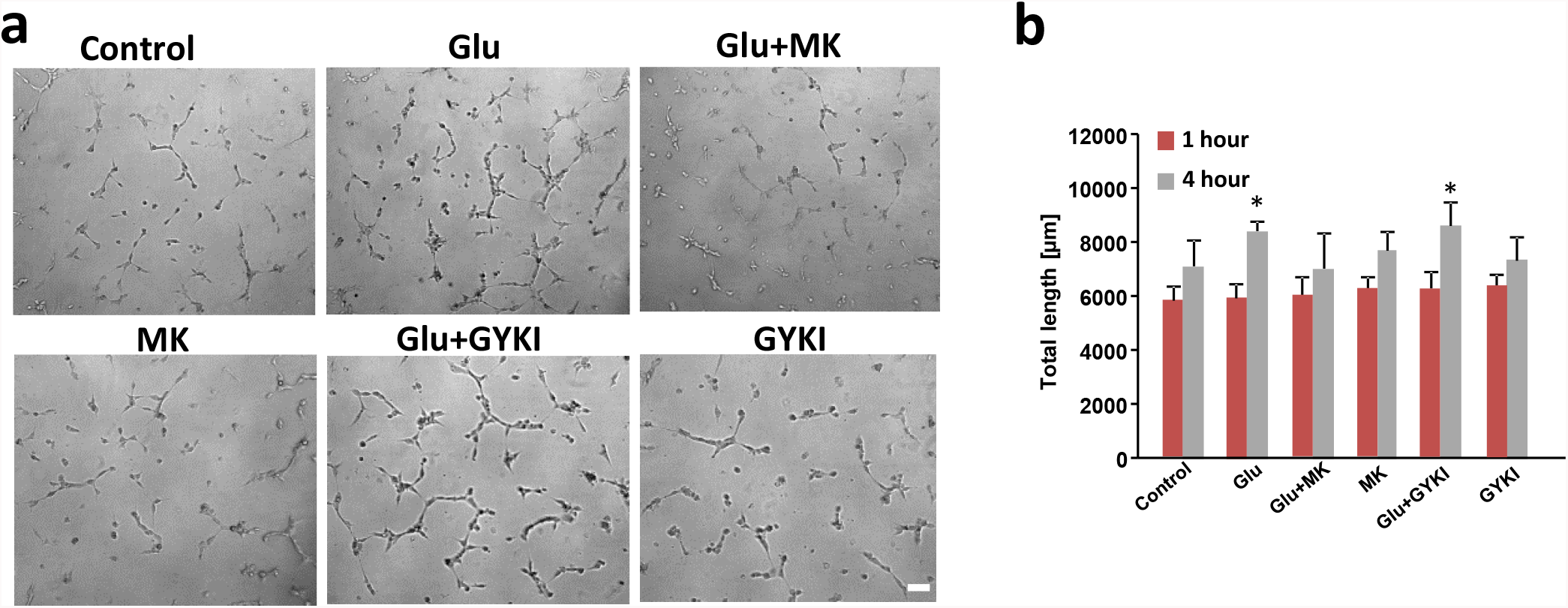
Glutamate promotes tube formation in brain endothelial cells. **a**, Glutamate promotes human umbilical vein endothelial cell (HUVEC) tube formation *in vitro*. Representative images show tube formation of endothelial cells under control conditions (control), after glutamate application (Glu, 100 µM), and with glutamate plus glutamate receptor antagonists MK801 (MK, 100 µM) and GYKI (50 µM). HUVEC cells were plated on Matrigel-coated plates and incubated solely in medium for 1 hour, images were taken and thereafter, endothelial cells were treated with PBS (control), glutamate and GluR antagonists for another 3 hours before tube formation was monitored again. HUVECs show enhanced tube formation capacity after glutamate treatment compared to controls while in MK+Glu treated group, the glutamate induced tube formation was blocked. GYKI application revealed no inhibiting effects on endothelial tube formation. MK801 or GYKI alone shows no endothelial toxic effect compared with control. Scale bar, 200 µm. **b**, Quantitative analysis of the total length of tubes in 1 hour and 4 hour time points. (n = 12 per group).

**Supplementary Figure 4:**
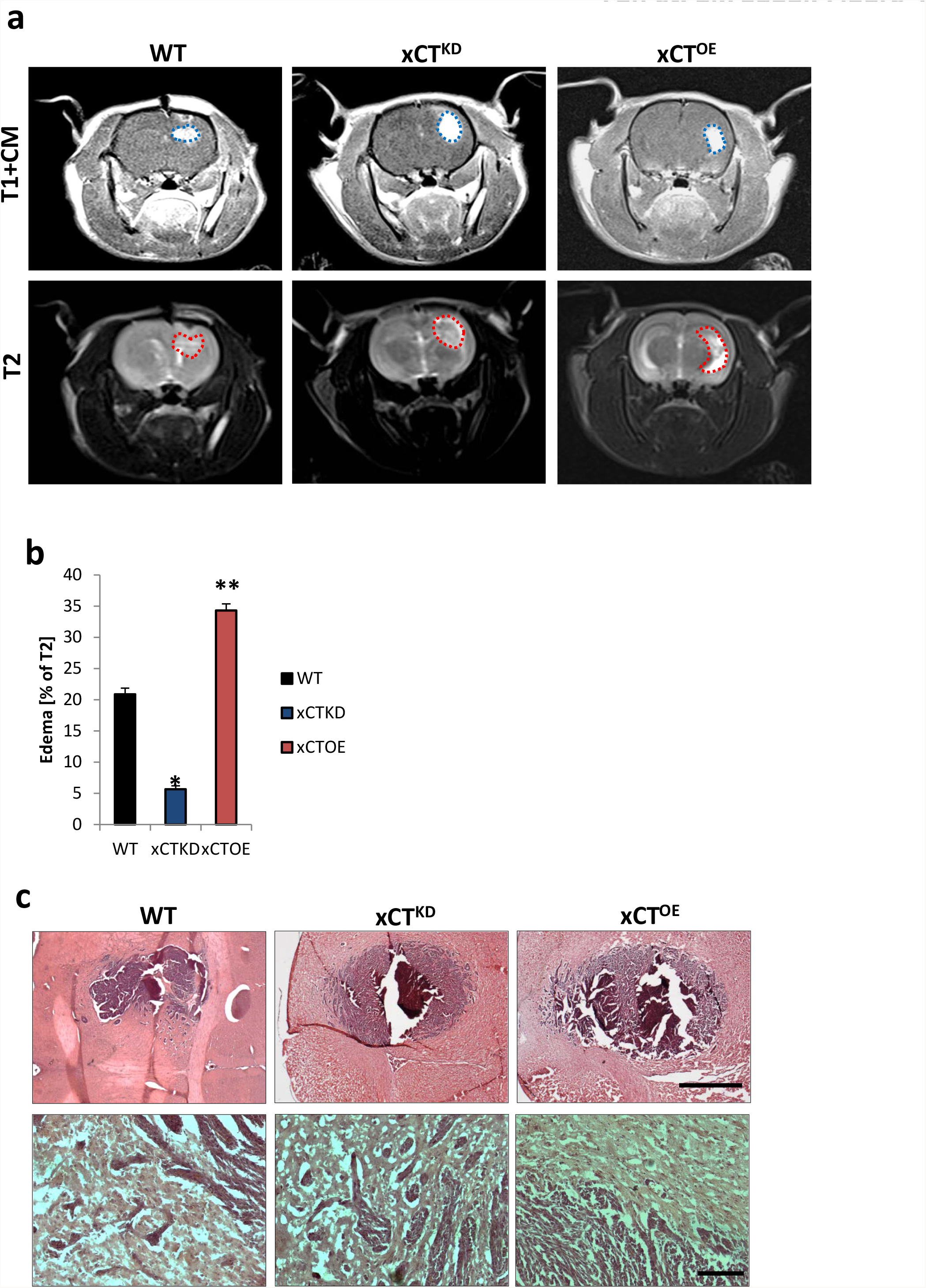
Enhanced xCT expression in gliomas causes increased brain edema. **a**, Representative MR images of wild-type (WT), xCT knockdown (xCT^KD^) and xCT-overexpressing (xCT^OE^) gliomas 10 days after tumor implantation in male Fisher rats weighing approximately 180–200 g. The tumor bulk (marked with blue dashed line) was visualized after application of intraperitoneal contrast agent and subsequently T1-weighted imaging (upper row, T1+CM). Bottom, corresponding T2-weighted images of brains from wild-type (WT), xCT knockdown (xCT^KD^) and xCT-overexpressing (xCT^OE^) gliomas. The red dashed line marked area indicates total tumor volume (including peritumor and edema zone). **b**, Quantification of edema volume out of T1 and T2-weighted MR images from wild-type (WT), xCT knockdown (xCT^KD^) and xCT-overexpressing (xCT^OE^) gliomas. Peritumor edema zone is significantly smaller in xCT knockdown gliomas compared to animals bearing wild type gliomas, while edema zone in xCT-overexpressing gliomas was significantly larger than wild type control group. (*P< 0.05, **P < 0.01). **c**, *Top*, Representative histological sections stained for HE from wild-type gliomas (WT), xCT knockdown (xCT^KD^) and xCT-overexpressing (xCT^OE^) gliomas. Scale bar represents 500 μm. *Bottom*, higher magnification in the area of tumor border. Scale bar represents 50 μm.

